# Whole Genome Doubling mitigates Muller’s Ratchet in Cancer Evolution

**DOI:** 10.1101/513457

**Authors:** Saioa López, Emilia Lim, Ariana Huebner, Michelle Dietzen, Thanos Mourikis, Thomas B.K. Watkins, Andrew Rowan, Sally M. Dewhurst, Nicolai J. Birkbak, Gareth A. Wilson, Mariam Jamal-Hanjani, Charles Swanton, on behalf of TRACERx Consortium, Nicholas McGranahan

**Author notes:** joint corresponding authors, or.

## Abstract

Whole genome doubling (WGD) is a prevalent macro-evolutionary event in cancer, involving a doubling of the entire chromosome complement. However, despite its prevalence and clinical prognostic relevance, the evolutionary selection pressures for WGD have not been investigated. Here, we explored whether WGD may act to mitigate the irreversible, inexorable ratchet-like, accumulation of deleterious mutations in essential genes. Utilizing 1050 tumor regions from 816 non-small cell lung cancers (NSCLC), we temporally dissect mutations to determine their temporal acquisition in relation to WGD. We find evidence for strong negative selection against homozygous loss of essential cancer genes prior to WGD. However, mutations in essential genes occurring after duplication were not subject to significant negative selection, consistent with WGD providing a buffering effect, decreasing the likelihood of homozygous loss. Finally, we demonstrate that loss of heterozygosity and temporal dissection of mutations can be exploited to identify signals of positive selection in lung, breast, colorectal cancer and other cancer types, enabling the elucidation of novel tumour suppressor genes and a deeper characterization of known cancer genes.

## Introduction

Whole genome doubling (WGD), involving the duplication of a complete set of chromosomes, is a common feature of cancer genomes (Zack, 2013, Bielski et al., 2018). It is thought to be induced through different mechanisms, including cytokinesis failure at the end of mitosis or endoreduplication, or more rarely, cell fusion (Davoli & de Lange, 2011), and has been linked to increased tumour cell diversity, accelerated cancer genome evolution and worse prognosis (Storchova & Pellman, 2004; Dewhurst et al., 2014, Bielski et al., 2018). Although WGD may lead to aneuploidy and chromosomal instability, it has been speculated that it may buffer the effects of chromosome or gene losses and deleterious mutations in tumor cells, favoring faster adaptive changes (Otto, 2007, Coward & Harding, 2014), analogous to the fitness effects resulting from polyploidy in other organisms.

Polyploidy is found across several plants and animal species in nature, and its evolutionary and biological significance has long intrigued researchers. Many hypotheses have been proposed about the selective advantages and/or disadvantages of polyploidy (Huxley, 1942; Madlung, 2013), but for some asexual species, like certain fungi (Selmecki et al., 2015) or plant lineages (Godfree et al., 2017), it has been demonstrated that ploidy variation could have a positive effect on fitness.

Specifically, it has been proposed that polyploidy could be a mechanism to overcome the Muller’s ratchet effect (Muller, 1964). Muller’s ratchet was originally described for asexual populations, where due to the lack of recombination, deleterious mutations would accumulate in an irreversible manner (Figure 1A). This phenomenon has been observed in nature in organisms such as the diploid Amazon molly (Loewe & Lamatsch, 2008), the self-fertile worm *C. elegans* (Loewe & Cutter, 2008), asexual DNA-based microbes such as *Salmonella typhimurium* (Andersson & Hughes, 1996) and amoebae (Maciver, 2016). The phenomenon is also described in the haploid setting in the context of the evolution of the Y-Chromosome (Engelstädter, 2008) and propagation of mitochondrial DNA (Loewe, 2006) in humans. It has also been linked with frailty, senescence and morbidity (Govindaraju & Innan, 2018). In the evolutionary history of species, the Muller’s ratchet effect has been countered by sexual reproduction, allowing recombination and the maintenance of or an increase in the diversity at the population level (Fisher, 1930; Muller, 1932). In asexual organisms, however, there are other mechanisms to counter the ratchet effect, including polyploidy (Maciver, 2016), which favors the accumulation of genetic variation promoting adaptation and diversity, allowing gene conversion, providing a strategy against genetic drift in asexual organisms.

Cancer development can be considered analogous to asexual evolution and, as such, due to lack of recombination, cancer cells may be subject to Muller’s ratchet, in particular in genomic segments with a high mutation rate exhibiting loss of heterozygosity (LOH) (Figure 1B).

Here, we explore the extent to which Muller’s ratchet can be observed in cancer genomes and consider whether a whole-genome doubling (WGD) event may act to buffer its deleterious impact. We focus on non-small cell lung cancer (NSCLC), the leading cause of cancer deaths worldwide (Bray et al., 2018), and one of the cancer types with the highest fraction of WGD in advanced tumours (Bielski et al., 2018). We explore this in the TRACERx (Tracking Non-Small-Cell Lung Cancer Evolution through Therapy) cohort (Jamal-Hanjani et al., 2017), a prospective and longitudinal study with multiregional data, and use the lung squamous cell carcinoma (LUSC) and lung adenocarcinoma (LUAD) data from the The Cancer Genome Atlas (TCGA) repository (Campbell et al., 2016) as a validation cohort. Furthermore, we also investigate how genome duplication can be exploited for identification of novel cancer genes and new therapeutic targets, and apply this approach to 33 different cancer types.

## Results

### Genome doubling has a marked positive impact on the viability of cancer cells subject to detrimental LOH

To explore the impact of a WGD event on cancer cell viability and fitness we first created a simplified model of cancer evolution, whereby cancer cells can gain fitness either through activation of oncogenes or inactivation of both copies of tumour suppressor genes, and, in order to incorporate a ratchet effect, we modelled the potential for loss of fitness through inactivation of essential genes (see Methods, Figure 1C,D). For simplicity, we do not incorporate a birth rate and only quantify the fitness of each individual cell. Thus, in our model, each cancer cell will eventually succumb to Muller’s ratchet and our analysis provides a quantitative framework to evaluate the impact of WGD, mutation rate (Figure 1C), and LOH (Figure 1D) on time to cancer cell death.

In every scenario, a WGD was found to increase the time to death of the cancer cell - i.e. providing cancer cells with a longer lifespan (at the likely detriment of the patient). In the case of varying mutations rates with a fixed LOH proportion of the genome of 0.28 - the median value of LOH proportions observed across all NSCLC datasets (TRACERx and TCGA) - (Figure 1C), we observed that a WGD event at time 30 increased the time to death ~6-10x. This increase was extremely relevant for cells with high mutation rates, which showed a sharp decline in viability without a WGD, with a mean time to death of 85 [83-87 95%CI] generations, due to the Muller’s ratchet process resulting from the irreversible accumulation of mutations in the absence of WGD. A WGD event, on the other hand, enabled the cells to live for an average of 683 generations [675-690 95%CI].

We also investigated the effect of WGD in different scenarios with varying LOH proportions and a fixed mutation rate of 0.000075 (Figure 1D). We observed that the LOH proportion was inversely correlated with the time to death, as high LOH proportions increase the probability of having essential genes affected by mutation. As before, an early WGD event at time 30 increased the time to death in all cases, from ~5x in the case of the lowest LOH proportion (0.1), to ~12x in cells with the highest fraction of the genome affected by LOH (0.6). Notably, cells with the highest proportions of LOH (0.6) and a WGD event survived longer (654 generations [546-662 95%CI]) than cells with the lowest proportions of LOH (0.1) without WGD (377 generations [370-383 95%CI]).

Even when considering a sharp decline in viability (reduced to half) due to the potential fitness costs of WGD, the duplication event was still found to increase the time to death between ~4-9x when varying LOH proportions and between ~4-6x with varying mutation rates (Figure S1).

Taken together, this model, despite simplifying cancer evolution, illustrates the importance of studying the biological relevance of WGD in cancer cells as a macro-evolutionary mechanism that might serve to buffer the Muller’s ratchet effect of deleterious mutations or copy number loss events in essential genes.

**Figure 1.**
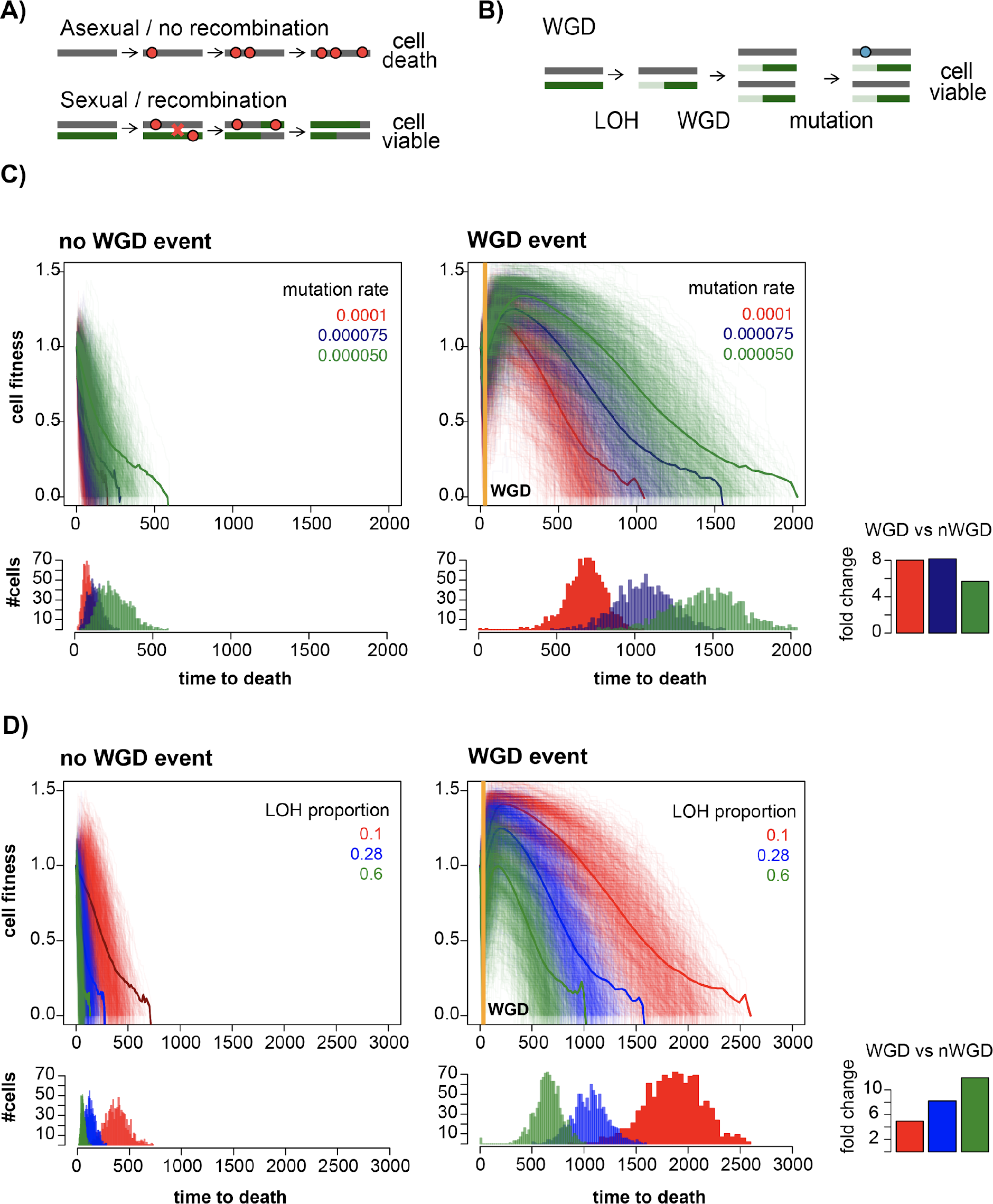
WGD buffers Muller’s ratchet. A) The principle of Muller’s ratchet in asexual and sexual organisms. The red dots represent the mutations acquired over time on chromosome segments. Asexual organisms with no recombination accumulate mutations in an irreversible manner leading towards cell death or extinction, while sexual populations with recombination are viable for longer periods of time. B) Muller’s ratchet in the context of cancer. WGD buffers the effects of late (post-WGD) deleterious mutations in regions of LOH by providing additional mutation-free segments. C) Simulations showing the effect of an early WGD event (orange line) on cancer-cell viability over time with different mutation rates (0.0001, 0.000075 and 0.000050) and LOH of 0.28. D) Simulations showing the effect of an early WGD event (orange line) on cancer-cell viability over time with different proportions of the genome subject to LOH (0.1, 0.28 and 0.6) and mutation rate of 0.000075.

### LOH and genome duplication are common events in NSCLC

Key assumptions in this model include that there is a sufficient burden of clonal LOH and/or a sufficiently high mutation rate in tumours and, crucially, that there is evidence for negative selection of essential genes in haploid regions, to foster an evolutionary benefit following a WGD event.

To explore the extent to which our simple model may accurately recapitulate lung cancer evolution, we first analysed the occurrence of WGD, the extent of LOH and the mutation rates of NSCLC from the TRACERx (Jamal-Hanjani et al., 2017) and TCGA datasets (Campbell et al., 2016). Genome doubling was identified as a frequent event in NSCLC (64.45% in LUAD and 66.12% in LUSC tumors, averaged across the TRACERx and TCGA cohorts), consistent with previous work (Jamal-Hanjani et al., 2017, Bielski et al., 2018). In the case of the TRACERx dataset, where multiregion data was available, WGD is primarily an early event, with only a small proportion being subclonal (6.25% in LUSC and 1.63% in LUAD) as previously reported in Jamal-Hanjani et al. (2017) (Figure 2A,B).

**Figure 2.**
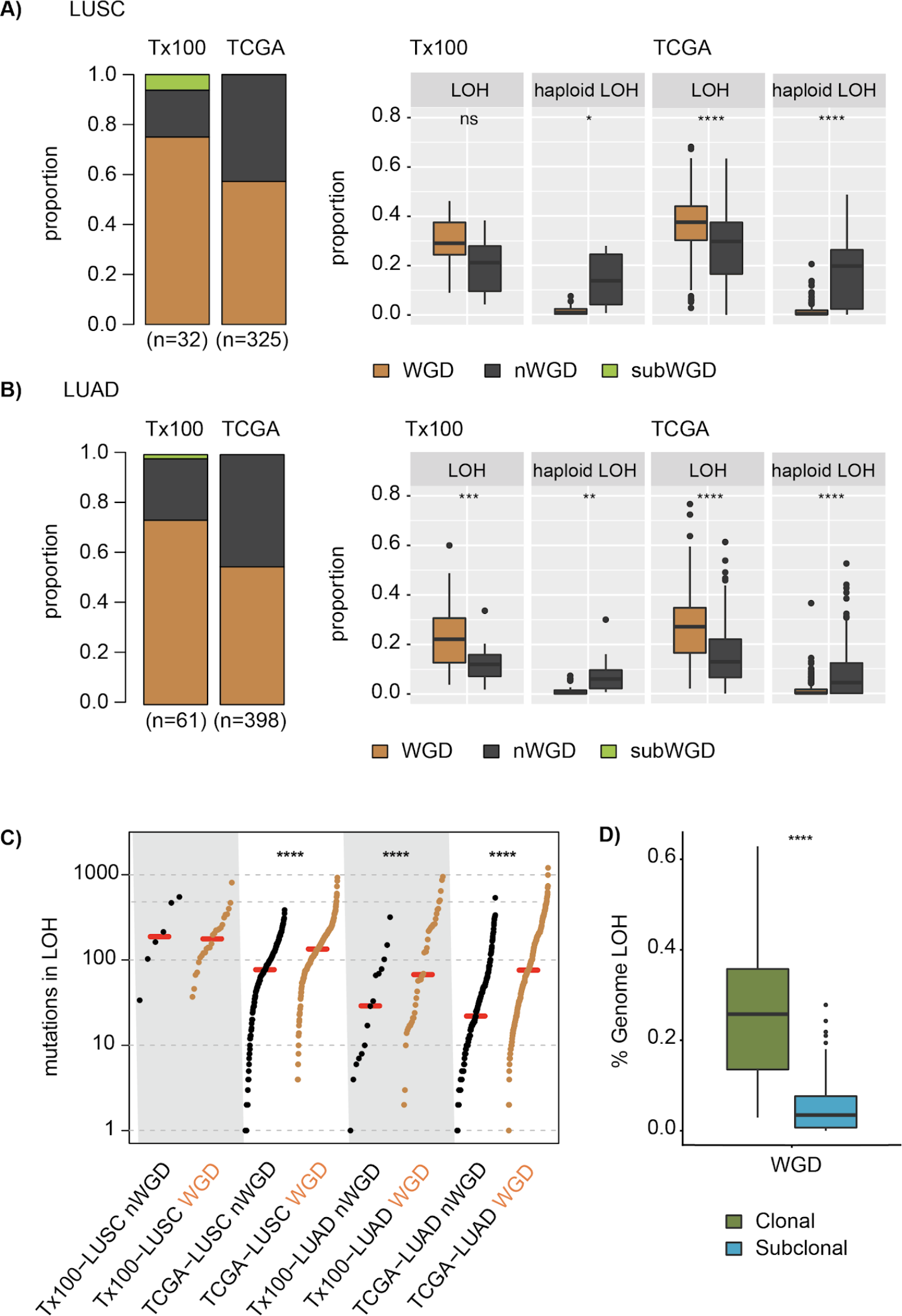
Prevalence of WGD and LOH in NSCLC. A-B) Proportion of WGD, subclonal WGD and non-WGD genomes (left), and proportion of the genome subject to LOH and haploid LOH in WGD vs nWGD (right) in LUSC (A) and LUAD (B). C) number of mutations in LOH, grouped by WGD status. The red lines indicate the median values. D) Differences in the proportion of clonal vs subclonal LOH in TRACERx data. Significant differences between the populations were assessed with a t-test. (* <0.05, ** p<0.01, *** p<0.001, **** p<0.0001)

We noted a significantly lower proportion of the genome subject to LOH in non-WGD tumours compared to their genome doubled counterparts (Figure 2A-C), but a higher proportion when looking at haploid LOH, as expected. Indeed, on average (across both cancer types and datasets) only 1.32% of the genome exhibited haploid LOH in WGD tumours, compared to 9.55% in nWGD, consistent with WGD reducing the haploid content of cancer cells (Figure 2A,B).

A fundamental assumption in our hypothesis is that LOH is predominantly a clonal event, occurring prior to WGD. Indeed, if LOH were to primarily occur after WGD, then the potential buffering effect of the duplication event would be negated, and LOH would frequently occur as a haploid event. We explored this matter by taking advantage of the TRACERx multi-region data. Figure 2D shows that the proportion of clonal LOH is significantly higher than the proportion of the genome displaying subclonal LOH (average of 25.85% vs 5.96%), suggesting the majority of LOH occurs early, and that GD tends to occur after a substantial amount of LOH is acquired, supporting the results from the model (Figure 1C,D).

### Accurate timing of mutations occurring pre and post WGD using HCT-116 cell lines

To quantify whether there is a shift in selection pressures following a duplication event, it is imperative to be able to time mutations - whether these occur before or after doubling. Previous work has suggested WGD provides a natural mechanism to temporally dissect mutations (Nik-Zainal et al., 2012; McGranahan et al., 2015). In brief, it has been assumed mutations occurring prior to WGD should be present at multiple copies, while those occurring after a doubling event would only be present at one copy (Figure 3A).

However, crucially, this assumption has not been subject to experimental validation. Therefore, we utilized an isogenic genome doubling model system involving genome-doubled HTC-116 clones deriving from non-genome doubled common ancestor (Dewhurst et al., 2014) and sequenced the whole-exomes of the ancestor plus two diploid and four tetraploid cell lines at two different time points (passages 4 and 50). Given that the shared common ancestor was diploid, all clonal mutations should be pre WGD, while private mutations should almost all be post WGD.

Reassuringly, we found 90.19% of pre-genome doubled mutations were correctly timed as such. Conversely, tetraploid private mutations, i.e. those that are absent in the diploid clones were correctly classified as late in 94.70% of the cases (Figure 3B). These data suggest that the inferences based on our timing approach are accurate and we can confidently distinguish which mutations occur before and which mutations occur after WGD. Applying temporal dissection of mutations to tumours exhibiting WGD in TRACERx and TCGA datasets suggests that the majority of detectable mutations in NSCLC accumulate prior to WGD (Figure 3C).

**Figure 3:**
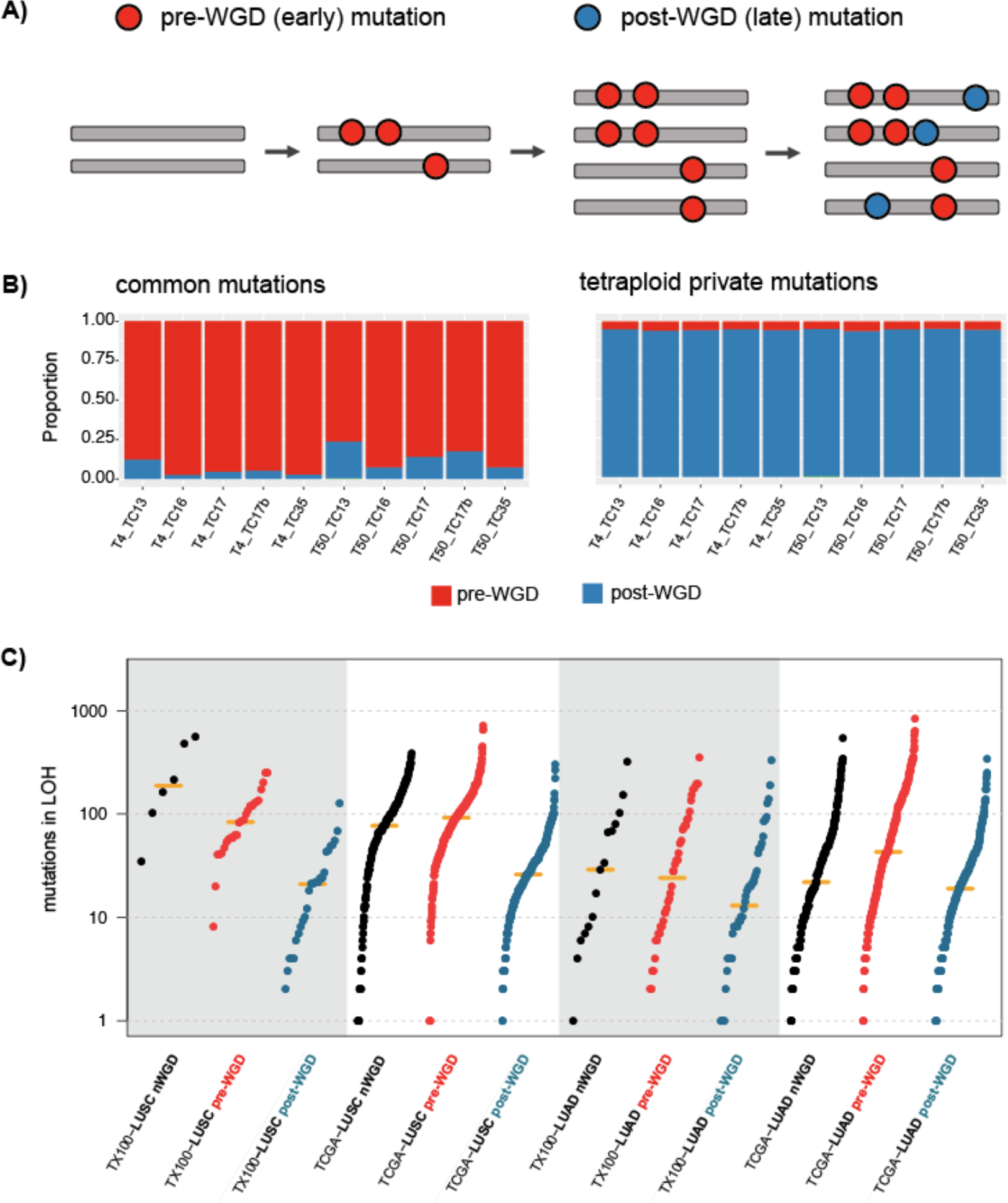
Timing mutations relative to WGD. A) Rationale of the mutation timing approach based on the copy number of mutations. Those mutations occurring before genome duplication (red dots) will be present at multiple copies, whereas those occurring after the duplication event (blue stars) will only be present at only one copy. B) Validation of the mutation timing approach using an isogenic genome doubling system involving genome-doubled HTC-116 clones deriving from a non-WGD common ancestor. Barplots show the proportion of mutations classified as either early or late by our timing approach for i) common mutations in the tetraploids (left) and ii) tetraploid private mutations (i.e., not present in the diploid genomes) (right). C) number of mutations in LOH across LUSC and LUAD datasets, grouped by doubling status and timing.

### Purifying selection may be operating on essential genes prior to duplication in regions of LOH in lung cancer

If WGD has a significant impact upon selection and the evolutionary course of the disease, we reasoned one would expect to see a difference in the selection of mutations occurring prior to duplication compared to those after the doubling event. Specifically, we hypothesized that purifying selection will operate on clones bearing deleterious mutations in housekeeping or essential genes in haploid regions of the genome, and such selection pressures will be relieved after genome duplication.

To investigate the selective pressures acting before and after WGD, and the effect of LOH, we applied a modified dNdS ratio to early (pre-WGD) and late (post-WGD) mutations within segments of LOH where we would expect evidence of purifying selection to be most profound (Martincorena et al., 2017). This dNdS ratio calculation assesses the fraction of non-synonymous mutations per non-synonymous site to synonymous mutations per synonymous sites, taking into account the propensity for different mutational processes to result in different mutation types. Under the assumption that synonymous mutations are neutral, this ratio can be informative about the direction of selection: ratios significantly above 1 are indicative of positive selection, while ratios below 1 are consistent with negative or purifying selection. As the selection signatures would vary depending on the biological functions of the genes in the evolution of cancer (Martincorena et al., 2017) we investigated different sets of genes, including essential genes identified by using extensive mutagenesis in haploid human cells (Blomen et al., 2015) and LUSC and LUAD-specific cancer genes described in the literature (Lawrence et al., 2014, Berger et al., 2016, Martincorena et al., 2017, Bertrand et al., 2018, Bailey et al., 2018). For our purposes, we looked at the dNdS values for truncating mutations (including nonsense and splice site mutations), which would be most closely associated with protein dysfunction and would in theory be subject to stronger purifying selection. dNdS values for missense mutations are shown in Figure S3.

Analyses at the whole exome level (all genes) showed no deviations from neutrality (Figure 4), as previously reported (Martincorena et al., 2017). However, consistent with a Muller’s Ratchet effect in NSCLC evolution, strong signatures of purifying or negative selection were observed for essential genes occurring in early mutations (pre-WGD) in genomic segments of LOH (Figure 4). These data suggest homozygous disruption to essential genes in regions of LOH in diploid genomes resulting in disruption of both alleles, leads to marked fitness impairment; indeed, on average, approximately three quarters of clones harbouring early truncating mutations in regions of LOH are predicted to be lost due to purifying selection.

**Figure 4.**
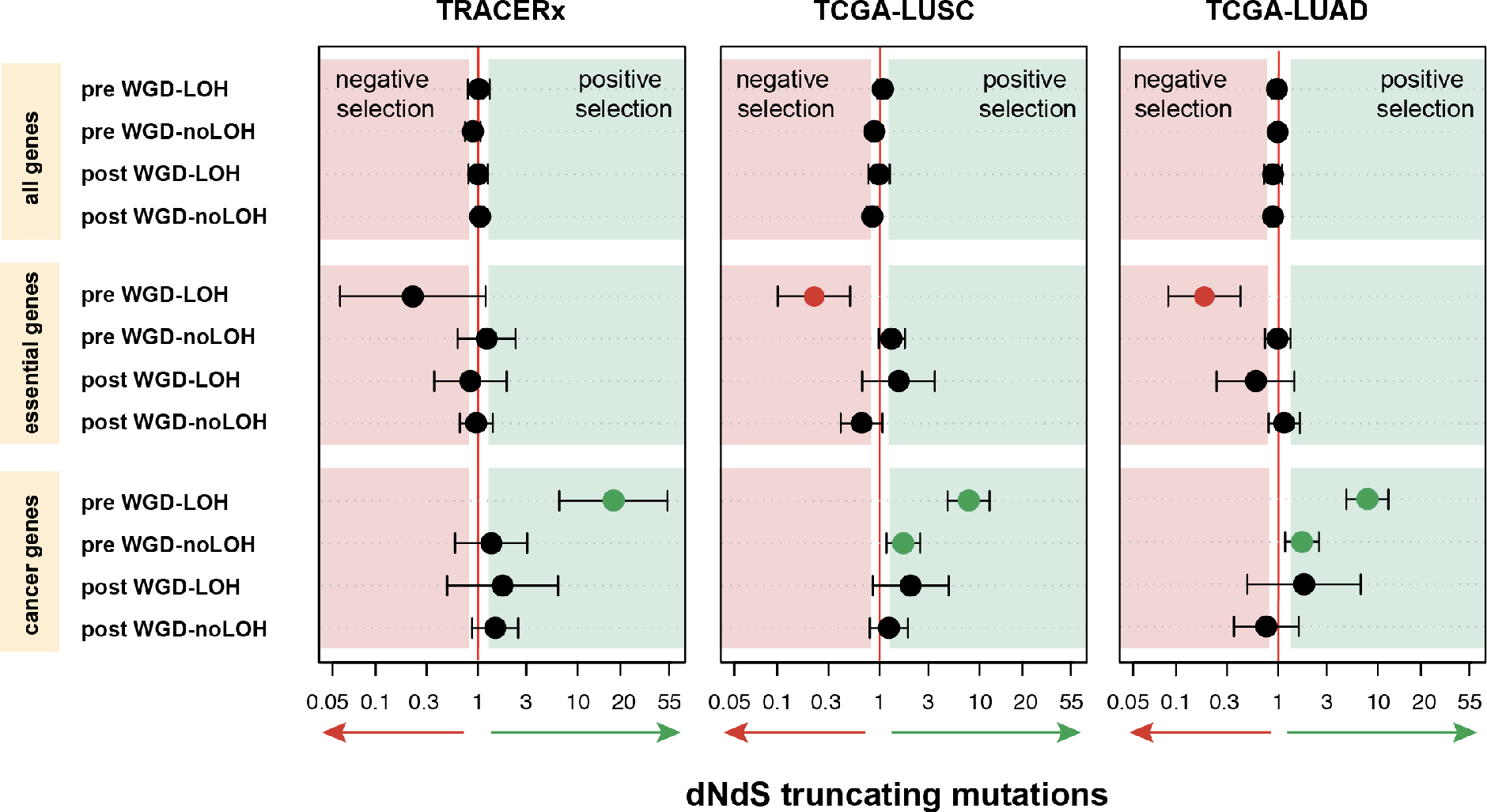
Purifying selection operates before but not after WGD. dNdS values for truncating mutations in WGD tumors calculated for all genes, essential genes and lung-specific cancer genes, grouped by LOH status and timing of the mutations in A) LUSC and LUAD from the TRACERx dataset (n=93), B) LUSC from TCGA (n=325) and, C) LUAD from TCGA (n=398). A dNdS ratio of 1 (red line) is consistent with neutrality. Values significantly higher than 1 (consistent with positive selection) are shown in green. Values significantly lower than 1 (indicating purifying selection) are shown in red. Notably, pre WGD mutations in LOH in essential genes are subject to purifying selection.

We hypothesised that following a WGD event (post-WGD), new mutations in genomic segments of LOH occur less frequently in a haploid context and therefore purifying selection of mutations in essential genes would not be observed. Consistent with this hypothesis, mutations occurring after WGD in essential genes were not found to be subject to significant purifying selection, with dNdS confidence intervals overlapping 1.

Likewise, in non-LOH regions, dNdS values were not significantly different from 1. These results suggest that genomic segments that accumulate truncating or nonsense mutations but still maintain one or more wild-type copies of the gene are not subject to strong negative selection as they are still viable. Therefore, by reducing the haploid genome fraction of cancer cells, WGD may act to mitigate the detrimental Muller’s Ratchet effect.

To rule out an excess of SNP contamination in essential genes leading to spurious signatures of negative selection we compared the proportion of somatic mutations contained in the SNP database (dbSNP) in essential (Blomen et al., 2015) vs non-essential genes and all genes (Figure S2). The proportion of mutations classified as SNPs was low in essential, non-essential and all genes in both LUAD (0.017, 0.022 and 0.022, respectively) and LUSC (0.025, 0.026 and 0.025, respectively), suggesting germline contamination of cancer exome reads is not responsible for the observed negative selection.

Essential genes can differ across cell and cancer types in a biologically meaningful manner. In order to assess the effect of using different essential gene lists, in addition to the essential genes from Blomen et al. (2015) identified by using extensive mutagenesis in haploid human cells (Figure 4), we quantified selection in other lists of essential genes from other works obtained through CRISPR-based systems (Hart et al., 2015, Wang et al., 2015). The results were highly consistent for TCGA-LUSC across the three different lists (Figure S4). For TCGA-LUAD, however, we note that the results are less consistent, suggesting that these alternative gene lists, obtained through CRISPR-based systems in various cell lines may lead to different results, potentially due to some genes acting in a haploinsufficient manner and others not.

Taken together, these data suggest there is strong negative selection against homozygous loss of essential genes in lung cancer. However, a genome duplication event can buffer the deleterious impact of a high mutation rate with high levels of LOH.

### Similar selection patterns are also observed across other cancer types

Prompted by these findings we sought to further investigate the Muller’s ratchet hypothesis across other cancer types from TCGA, which show a notable variability in the fractions of WGD, haploid LOH (Figure S5) and mutation rates. Notably, consistent selection patterns were observed among most of those cancer types with enough patients/mutations to perform selection analyses (Figure S6). Similar to NSCLC, significant signatures of purifying selection in pre-WGD truncating mutations in LOH in essential genes (but not after WGD) were observed in skin cutaneous melanoma (SKCM), characterized for having very high mutation rates. In other cancer types (liver hepatocellular carcinoma -LIHC-, colon adenocarcinoma -COAD-, uterine corpus endometrial carcinoma -UCEC-, bladder urothelial carcinoma -BLCA-, cervical squamous cell carcinoma and endocervical adenocarcinoma -CESC-, head and neck squamous cell carcinoma -HNSC-, esophageal carcinoma -ESCA-), although not significant, a similar trend was also observed. In the case of breast invasive carcinoma (BRCA) no evidence of negative selection acting on essential genes prior to WGD was observed, potentially due to low mutation rate observed in this cohort as a whole.

### Leveraging mutations with genomic segments in LOH to identify new driver genes across cancer types

Positive selection is a feature of cancer, reflected as an enrichment of non-silent mutations in cancer genes (Martincorena et al., 2017., Tamborero et al., 2013, Zapata et al., 2017), and thus the dNdS ratio can be exploited not only to identify signals of selection but also used to identify new driver genes in cancer. We therefore next explored dNdS values specifically for known cancer genes. We observed highly significant positive selection of truncating mutations prior to WGD in regions of LOH, followed by reduced positive selection after duplication (Figure 4). These data are consistent with evidence for strong selection for inactivation of both copies of tumour suppressor genes through LOH and mutation, prior to WGD.

We hypothesized therefore that by specifically looking into genomic segments of LOH we could identify novel tumour suppressor genes that would not be otherwise detected. Moreover, we could explore whether selection intensities varied depending on whether the mutation co-occurred with LOH or the wild type allele remained intact.

We obtained dNdS values for nonsense mutations at the gene level and compared the selection intensities for early mutations in regions of LOH (Figure 5A, x-axis) compared to mutations not occurring in segments of LOH (Figure 5A, y-axis). Figure 5A shows the genes that were significantly selected when looking solely at early mutations in LOH segments (blue rectangles), versus all regions (orange rectangles), or both (grey rectangles). Notably, two genes, with very high selection coefficients in LOH regions but lower in non-LOH regions in LUSC were *TP53* and *PTEN* (Figure 5A, left). These are well-established tumor suppressor genes that, concordant with Knudsen’s “two-hit hypothesis”, require disruption of both alleles for optimal fitness potential to be achieved. The fact that the selection coefficients are significantly higher when looking at mutations in LOH supports this notion, and importantly, it also suggests that this approach could be used to identify novel tumor suppressor genes in cancer genomes. This could be the case of *ZNF750* (Zinc Finger Protein 750), subject to high positive selection in regions of LOH in LUSC, and would have remained undetected with standard procedures looking at the whole mutation load. Intriguingly, *ZNF750* has previously been described as a lineage-specific tumour suppressor gene in squamous cell carcinoma (Hazawa, 2017), consistent with its occurrence as an early event in 87% (7/8) of LUSC tumours. Other genes that were only identified as significant when looking at mutations in regions of LOH were *NOTCH1* and *SMAD4* in LUSC. On the other hand, *CUL3* (Cullin-3), which plays an important role in the ubiquitin-proteasome system (Chen, 2016), showed high signatures of selection in regions not subject to LOH, possibly reflecting haploinsufficient activity in cancer evolution, and, conceivably, the requirement for an intact wild-type allele to reach an optimal fitness potential.

In the case of LUAD (Figure 5A, right), we identified two genes not currently considered as validated cancer-genes in COSMIC database: *CMTR2* (Cap Methyltransferase 2), also proposed as a LUAD cancer gene by Martincorena et al., 2017, and *MGA* (MAX dimerization protein), a transcription factor that has also been suggested as a tumor suppressor gene in LUAD previously (Jamal-Hanjani et al., 2017, Bailey et al., 2018).

As an orthogonal approach for the identification of driver genes under different scenarios (mutations in LOH, non-LOH, all) we also implemented MutSigCV (Lawrence et al., 2013). Figure S7 shows the q values obtained for each gene, coloured according to their presence in COSMIC database (red=not included, black=included). In both LUSC and LUAD, we detected potential driver genes when looking at LOH mutations that were not significant when assessing all mutations together, or mutations in non-LOH. These include *SERINC2*, *NPRL2*, *NCKIPSD*, *MEGF9*, *LRRC8E* and *ROR2* in LUSC and *ZBTB6*, *PHLDA1* and *FLYWCH2* in LUAD.

Finally, we extended the analyses to the remaining cancer histological subtypes in the TCGA cohort. Following the same approach as above, it was possible to identify a total of 27 additional potential tumour suppressor genes across cancer types by limiting the analyses to segments of LOH (Figure 5B). Some of these are included in the COSMIC database as well-characterized cancer genes in some cancer types (*RB1*, *PTEN*, *SMAD4*, *BAP1*, *SETD2*, *NOTCH1*, *CDH1*, *BRCA1*, *NF2*, *SMAD3*, *PTCH1*, *ITGAV* and *CDK12*). However, we also identified additional genes under positive selection not included in the current release of COSMIC: *ZNF750*, *ARHGAP35*, *WWC1*, *TUBA3C*, *NCLN*, *KRTAP19-5*, *GRIK2*, *GLRA1*, *AFXDC2*, *FAM19A3*, *CRYGC*, *CLEC4E*, *CCAR1* and *AC061992.1*.

**Figure 5.**
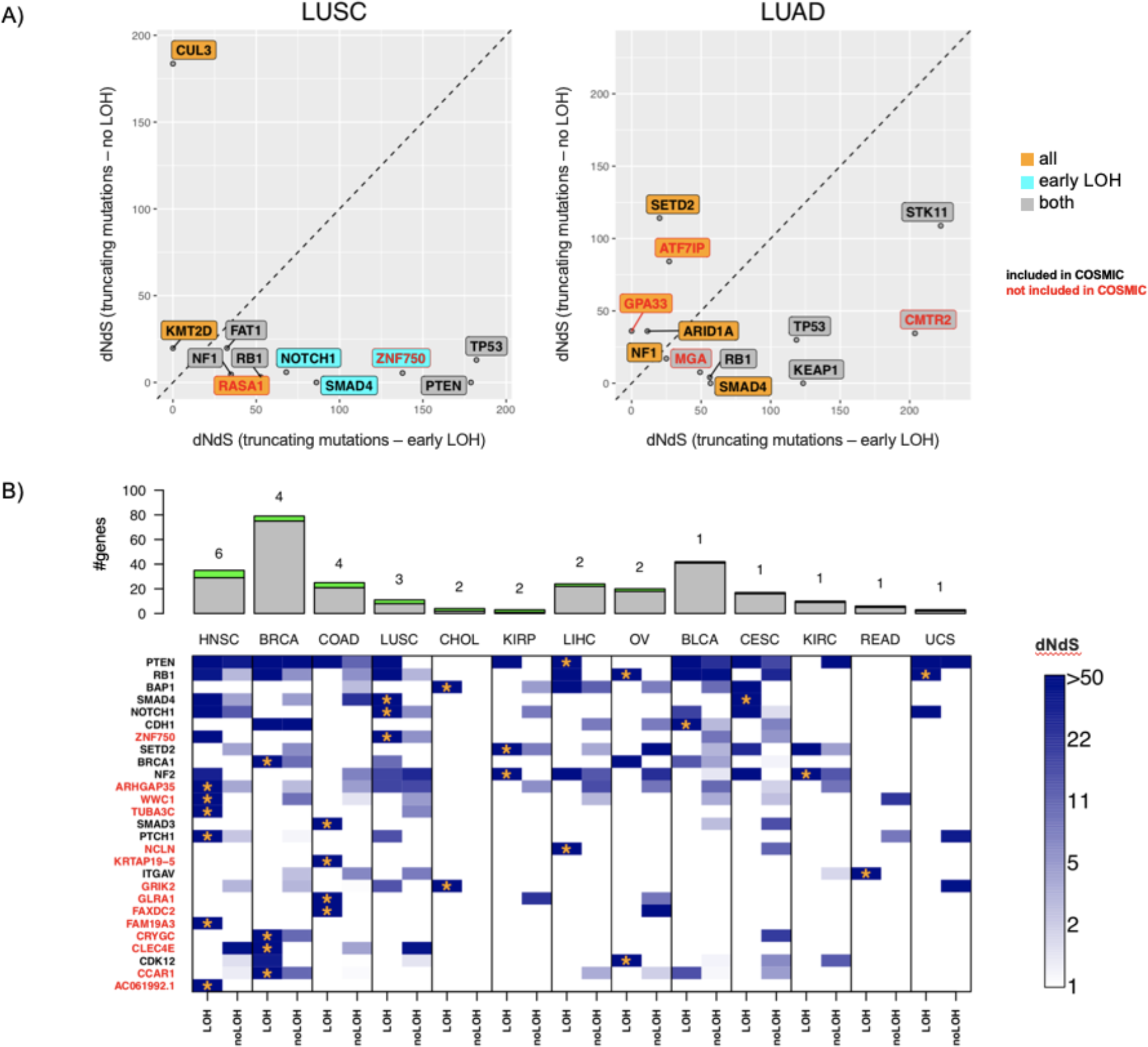
Exploiting LOH to identify cancer genes. A) dNdS selection coefficients for truncating mutations in early mutations in LOH (x-axis) vs truncating mutations in genomic regions without evidence of LOH (y-axis). The background color indicates whether the gene was identified as significant, using mutations in early LOH (cyan blue) using all mutations (yellow), or identified as significant in both cases (gray) (q-value<0.05). The text color represents whether the gene is currently included in the COSMIC database as a cancer gene. B) dNdS selection coefficients for truncating mutations in early mutations in LOH (“LOH”) vs truncating mutations in regions without evidence of LOH (“noLOH”) across cancer types. Only genes that are significant in at least one cancer type in the LOH category are shown. Barplots show the total number of cancer genes that are significantly (q-value<0.05) identified using the “all approach” (grey) and the number of cancer genes that are only identified (q-value<0.05) using the “LOH” approach (green). The latter is also represented with a number above the bars, and the specific genes are marked with orange stars in the heatmap. Only those cancers where we identify additional cancer genes using the “LOH” approach (this is, only looking at early mutations in LOH) are shown. (HNSC=head and neck squamous cell carcinoma, BRCA=breast adenocarcinoma, COAD=colon adenocarcinoma, LUSC=lung squamous cell carcinoma, CHOL=cholangiocarcinoma, KIRP=kidney renal papillary cell carcinoma, LIHC=liver hepatocellular carcinoma, OV=ovarian serous cystadenocarcinoma, BLCA=bladder urothelial carcinoma, CESC=cervical squamous cell carcinoma and endocervical adenocarcinoma, KIRC=kidney renal clear cell carcinoma, READ=rectum adenocarcinoma, UCS=uterine carcinosarcoma).

## Discussion

Despite the fact that WGD is associated with poor prognosis and an acceleration of chromosomal instability, a rational basis for the observation of the recurrence of WGD events in human tumors remain unclear. Cancer progression is an evolutionary process, and as such, the fundamental principles of Darwinian evolution and modelling of ecological systems can be applied to study tumor development. In this work, we utilize concepts from species evolution to shed light on the evolutionary importance of WGD in cancer, and reveal how this can be exploited to reveal novel cancer genes.

While human germline evolution is dominated by a relatively low mutation rate and negative or purifying selection, which removes deleterious or harmful mutations, cancer evolution is characterized by high mutation rates and pervasive positive selection (Martincorena et al., 2017). Indeed, the extent of negative selection during somatic evolution has been subject to debate. For example, while negative selection has been reported in transcription factor binding motifs (Vorontsov et al., 2016), hemizygous regions (van den Eynden et al., 2016; Martincorena et al., 2017) and splicing-associated sequences (Hurst et al., 2017), conflicting reports regarding the extent to which the immune system results in negative selection and mutation loss have been presented (Zapata et al., 2018; Martincorena et al., 2017; Van den Eynden et al., 2018).

In this work, we dissected mutations temporally relative to WGD, permitting an exploration of whether selection pressures change following WGD, both in the context of essential genes and cancer genes. Focussing on genomic segments exhibiting LOH, we demonstrate that truncating mutations occurring before WGD in a haploid context in essential genes are subject to strong negative selection in lung cancer evolution. Analogous to haploid asexual and non-recombining populations in nature, cancer cells will accumulate these mutations irreversibly, in a ratchet-like process. Thus, in cancers with high mutation rate and levels of LOH, and in the absence of other compensating mechanisms, this may lead to the attrition of subclones.

Mechanisms to counteract the ratchet-like process of mutation acquisition in haploid genomic regions may therefore be beneficial; indeed, in species evolution, sexual reproduction and polyploidy have been postulated to fulfil these roles (Muller, 1964). Using a dNdS selection analysis (Martincorena et al., 2017), we find that an early WGD, may eliminate the requirement for purifying selection. Specifically, by duplicating haploid genomic segments (which hence become diploid), WGD may drastically attenuate cancer cell attrition through disruption of both copies of essential genes. This is also supported by the observation that LOH is a clonal phenomenon in NSCLC progression that predates the duplication of the genomes, in line with recent work by Bielski et al. (2018), supporting a model by which WGD occurs early in cancer evolution. Consistent with this, our simplified model of cancer evolution suggests an early WGD is associated with a ~10x increase in lifespan of cancer cells (Figure 1C,D).

A more sophisticated model may shed light on the tipping point at which levels of LOH combined with tumor mutation burden in a tumor would lead to a WGD being positively selected. Such a model may shed light on the divergent frequencies of WGD events across cancer types. Indeed, our model only likely applies to cancer types with a sufficiently high mutation rate and LOH proportion. While this is likely the case in NSCLC, as shown in this work, it can also be applied to other cancer types like melanoma (SKCM), where the same selection patterns are observed. However, there are other cancer types, like testicular germ cell tumors (TGCT) which, despite having low mutation rates, display high WGD rates. In these cases the underlying reasons for the existence of WGD cannot be explained by the Ratchet effect, highlighting that the countering the Muller’s Ratchet is likely not the sole reason for WGD.

Conceivably, in the context of treatment that induces hypermutation, WGD may also be selected, particularly in the context of tumours with high LOH. For example, Temozolomide (TMZ) is an alkylating agent used to induce apoptosis in glioma, melanoma and other cancer types. This agent is also mutagenic, and it has been shown that the introduction of thousands of *de novo* mutations may drive the evolution of TMZ-resistant tumor cells to higher states of malignant potential (Johnson et al., 2014). Speculatively, WGD may buffer the deleterious impact of therapy-induced mutations, thereby facilitating treatment resistance.

An additional contribution from this work is the identification of 27 novel potential lung cancer driver genes, identified by considering genes within segments of LOH. Temporal dissection of mutations, coupled with a focus on regions of the genome exhibiting LOH, enables elucidation of genes subject to two-hits (mutation and LOH) and strong signals of positive selection. Importantly, the signal of positive selection may be missed without such dissection. Indeed, although further experimental validation is required, our framework enabled elucidation of several putative cancer genes that are not currently included in the COSMIC database (including *GLRA1* in COAD, *CRYGC* in BRCA or *FAM19A3* in HNSC). In addition, our results confirm that many established tumour suppressor genes, including *PTEN* and *RB1*, likely require both hits to be subject to positive selection, while other cancer genes, including *CUL3*, are subject to strong positive selection without two hits. Conceivably, a similar framework could be applied to identify cancer genes subject to either mutation and methylation, or methylation and LOH.

In conclusion, our study highlights the parallels between species and cancer evolution and emphasizes the importance of macro-evolutionary events in cancer development.

## Methods

### Simulations

We performed simulations to model the viability of a cancerous cell over time and illustrate the potential of increasing DNA copies via genome duplication in the mitigation of damaging effects of mutations. This simplistic model assumes that each cell starts with a fitness of 1, varying LOH proportion in the genome (ranging from 0.1 to 0.6) and varying mutation rates (ranging from 0.00075 to 0.001). Each cell has a total of 30,000 genes, of which 1,400 are essential genes, 300 oncogenes and 300 tumor suppressor genes.

The viability of the cell varies by a function of the mutations that the cell has accumulated over time, which exert deleterious or beneficial effects on cell fitness to varying degrees, depending on the type of genes where they are accumulated. If mutations arise in essential genes, the viability decreases 0.2 units when there are no wild type copies of the genes left. On the contrary, if mutations arise in tumor suppressor genes with no wild type copies of the genes left, or in oncogenes, the viability increases 0.1 units. In each run, a maximum of 5 oncogenes and 5 tumor suppressor genes (the 2 alleles) are allowed to be mutated. Finally, if mutations are in other genes (non-essential, non-oncogenes, non-tumor suppressors), with no wild type allele left, the viability declines 0.1. In order to account for the possible fitness costs associated with genome duplication, we simulated alternative scenarios including a reduction in viability of 50% following the duplication event.

We simulate the process until viability reaches 0, indicative of cell death, and record the time. We performed 1000 iterations of each run and used a linear interpolation calculated with the approx() function in R to find the average curve.

### Data processing

We analyse 93 patients from the first cohort of patients of NSCLC (61 LUSC and 32 LUAD) analyzed by the lung TRACERx (TRAcking Cancer Evolution through therapy (Rx)) project and thoroughly described in Jamal-Hanjani et al. (2017).

Additionally, raw .bam files for LUAD (n=398) and LUSC (n=325) samples from the TCGA repository were downloaded and processed through the TRACERx pipeline. Briefly, we used BWA-MEM to align the reads to the reference genome (build hg19). We used Platypus for SNP calling on the germline, and Varscan2 and MuTect for somatic mutation calling. Functional annotation of genomic variants was performed using ANNOVAR (Wang K et al., 2010). Purity, ploidy and copy number profiles of tumor cells were obtained with ASCAT (Van Loo et al. 2010), using the matching germline data. Mutations were timed as early or late based on the mutation and major allele copy number. Following a conservative approach, we considered early mutations those with mutation copy number >=1.75 and major allele copy number >=1.75. Mutations were classified as late if mutation copy number <=1.25 and major allele copy number >=1.75. Clonal mutations that could not be timed were classified as “unknown”. Additionally, mutations were defined as mutations in LOH if the minor allele copy number was <0.25. The WGD status for each tumor was obtained using the genome doubling algorithm described in Dewhurst et al. (2014).

For the identification of new drivers across other non-lung cancer types, we downloaded the MC3 somatic mutation tables (Ellrott et al., 2018) from whole-exome sequencing data across 31 cancer types from TCGA. Further processing of the samples was performed as described above.

### Selection tests

Selection across LUSC and LUAD was quantified using the dNdScv R-package (Martincorena et al., 2017), an implementation of the traditional dNdS ratio, adapted and refined for cancer genomes. In all cases we used the trinucleotide substitution model with 192 rate parameters and default parameters, removing ultra-hypermutator samples (3000 maximum number of coding mutations per sample) and limiting the analyses to 3 mutations per gene per sample. dNdS ratios were quantified for missense, nonsense and essential splice mutations, both globally and at gene level.

Global dNdS values for nonsense mutations were calculated for different lists of genes: a) *all genes*, b) *cancer genes*, which includes a compilation of LUSC and LUAD-specific tumor suppressor genes and oncogenes described in the literature (Lawrence et al., 2014, Berger et al., 2016, Martincorena et al., 2017, Bertrand et al., 2018, Bailey et al., 2018) and c) *essential genes*, which includes the essential genes reported by Blomen et al., 2015, identified by using extensive mutagenesis in haploid human cells (1154 genes). Different lists of essential genes obtained through CRISPR-based systems were also evaluated for comparison (Hart et al., 2015, Wang et al., 2015). Essential genes from Hart et al. (2015) included the 1580 hits observed in three or more of the five cell lines. In the case of Wang et al., (2015), we included the genes with adjusted p-values <0.05 in three or more of the four cell lines used (a total of 1282 genes).

Selection tests were performed on different subsets of mutations considering the timing (pre-WGD vs post-WGD mutations) and the LOH presence (LOH vs nonLOH) in WGD genomes.

### Identification of driver genes

Maximum-likelihood dNdS estimates (MLE) at the gene level obtained with the dNdScv package were used to identify potential driver genes under positive selection. We compared the MLE values for nonsense mutations (wnon) in early LOH vs non-LOH.

As an additional method to identify driver genes we used MutSigCV (Lawrence et al., 2013), comparing the adjusted p-values for mutations in LOH, non-LOH and all mutations combined. In order to reduce the number of significant hits we filtered the genes that are expressed at low levels (below the median expression value across all the genes and patients) in that particular cancer type. RNAseq-based normalized expression data for each cancer type was obtained from the TCGA data portal.

We also sought to extend our LOH-based approach to other non-lung cancer types using the MC3 somatic mutations tables from TCGA database, as described in the Data processing section.

### Validation of the timing approach using mutation and copy number data from diploid and tetraploid clones from the HCT-116 cell line

In order to demonstrate that we can accurately quantify which mutations occur pre and post the genome duplication event we used mutation and copy number data from 2 diploid and 4 tetraploid subclones derived from diploid human colon carcinoma HCT-116 cells, isolated at different passages (4 and 50 passages) (Dewhurst et al., 2014).

For each sample exome capture was performed on 1-2 ug DNA isolated from genomic libraries with median insert size of 190bp, using a customised version of the Agilent Human All Exome V5 kit, according to the manufacturer’s protocol (Agilent). Samples were 100bp paired-end multiplex sequenced on the Illumina HiSeq 2500 at the Advanced Sequencing Facility at the Francis Crick Institute. The data was aligned to the reference human genome (build h19) and somatic mutations were then obtained with Varscan and MuTect, using the parental diploid clone as the germline. Copy number profiles were inferred with PICNIC using genotype chip data. The early (pre-WGD)/late (post-WGD) classification of the mutations, and purity and ploidy of the cell lines was assessed using the same procedure as above.

### Statistics

R version 3.3.1 was used to analyze the data. No statistical tests were used to predetermine the same size. Tests involving comparison of distributions were done using ‘t.test’ using the unpaired option, unless otherwise stated. Confidence intervals for dNdS analysis was obtained using the dNdSCV package (Martincorena et al., 2017).

### Data availability

Sequence data used during the study will be deposited at the European Genome-phenome Archive (EGA), which is hosted by The European Bioinformatics Institute (EBI) and the Centre for Genomic Regulation (CRG) under the accession code: EGAXX. Further information about EGA can be found at https://ega-archive.org.

## Acknowledgements

S.L. receives funding from Rosetrees.

C.S is Royal Society Napier Research Professor. C.S. is funded by Cancer Research UK (TRACERx and CRUK Cancer Immunotherapy Catalyst Network), the CRUK Lung Cancer Centre of Excellence, Stand Up 2 Cancer (SU2C), the Rosetrees and Stoneygate Trusts, NovoNordisk Foundation (ID 16584), the Breast Cancer Research Foundation (BCRF), the European Research Council (THESEUS), Marie Curie ITN Project PLOIDYNET (FP7-PEOPLE-2013, 607722), the NIHR BRC at University College London Hospitals, and the CRUK University College London Experimental Cancer Medicine Centre.

N.M is a Sir Henry Dale Fellow, jointly funded by the Wellcome Trust and the Royal Society (Grant Number 211179/Z/18/Z), and also receives funding from CRUK, Rosetrees, and the NIHR BRC at University College London Hospitals.

## Author contributions

Conceptualization and supervision: N.M and C.S. Manuscript preparation: S.L., N.M. Manuscript review/editing: S.L., C.S., N.M. Formal analysis: S.L., E.L. Visualization/data presentation: S.L. Data curation and interpretation of results: S.L., E.L., A.H., M.D., T.M., T.W., N.B., G.W., N.M. Resources: M.J.H, C.S, N.M.

