## Supplementary figures for "Whole Genome Doubling mitigates Muller’s Ratchet in Cancer Evolution"

**Figure S1.** Simulations showing the effect of an early WGD event on cell viability with varying A) LOH proportions and B) mutation rates. Here the WGD involves a high fitness cost associated with increased energy-demanding metabolic reactions, the replication of duplicated DNA and gene expression.

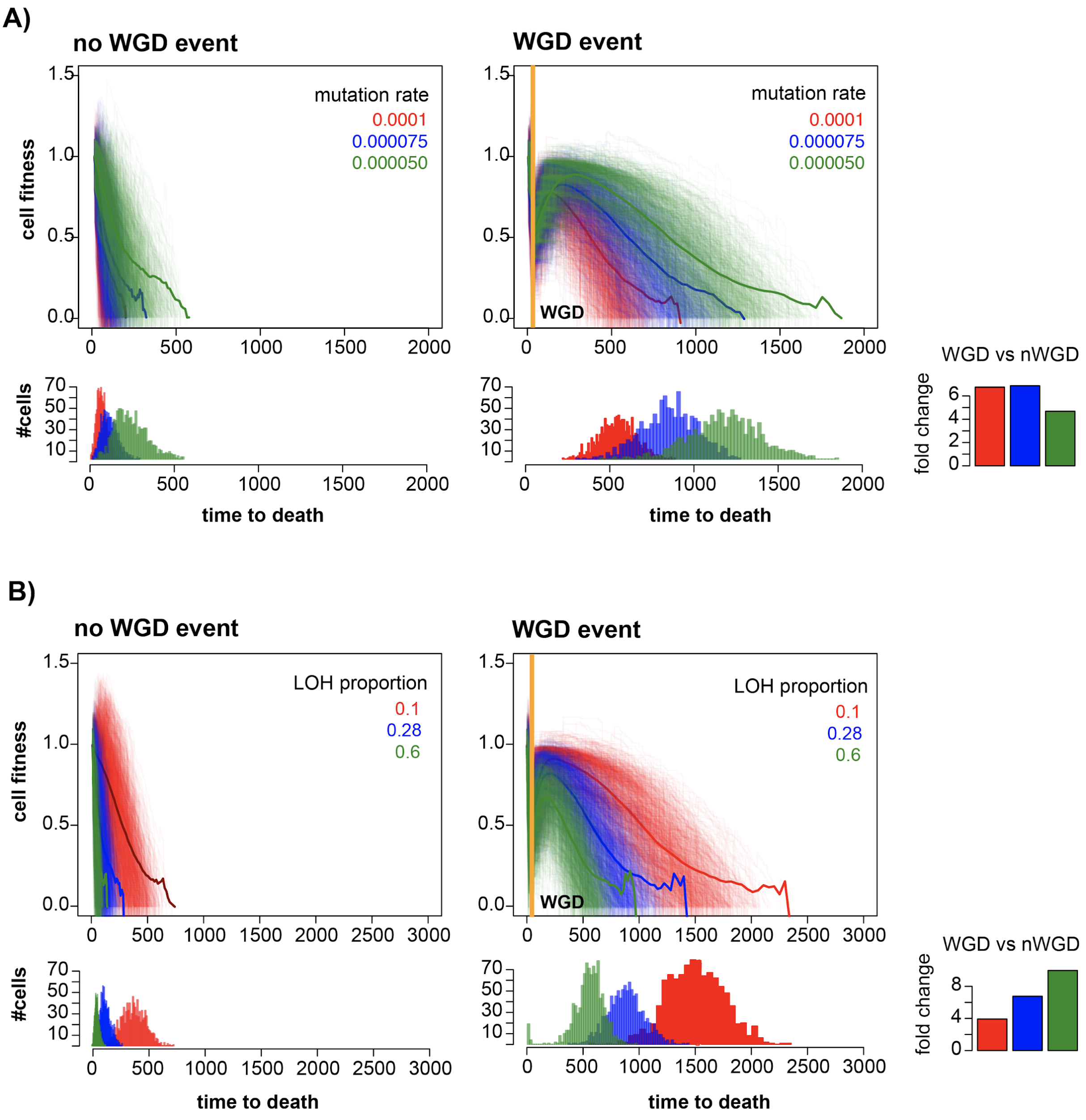

**Figure S2.** Proportion of dbSNPs included in the somatic data in all genes, essential (Bloomen et al., 2015) and non-essential genes in LUAD and LUSC.

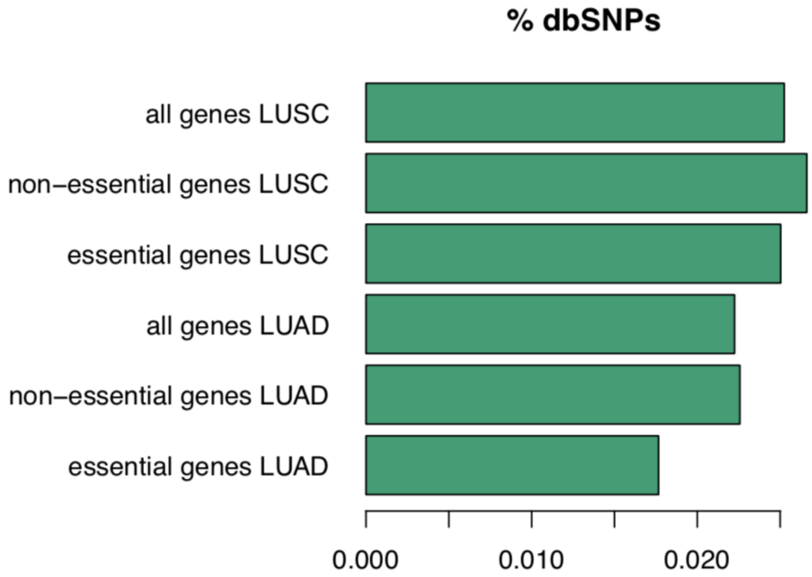

**Figure S3** - dNdS values for missense mutations in WGD tumors calculated for essential genes, lung-specific cancer genes, and all genes, grouped by LOH status and timing of the mutations in A) LUSC and LUAD from the TRACERx dataset (n=93), B) LUSC from TCGA (n=325) and, C) LUAD from TCGA (n=398).

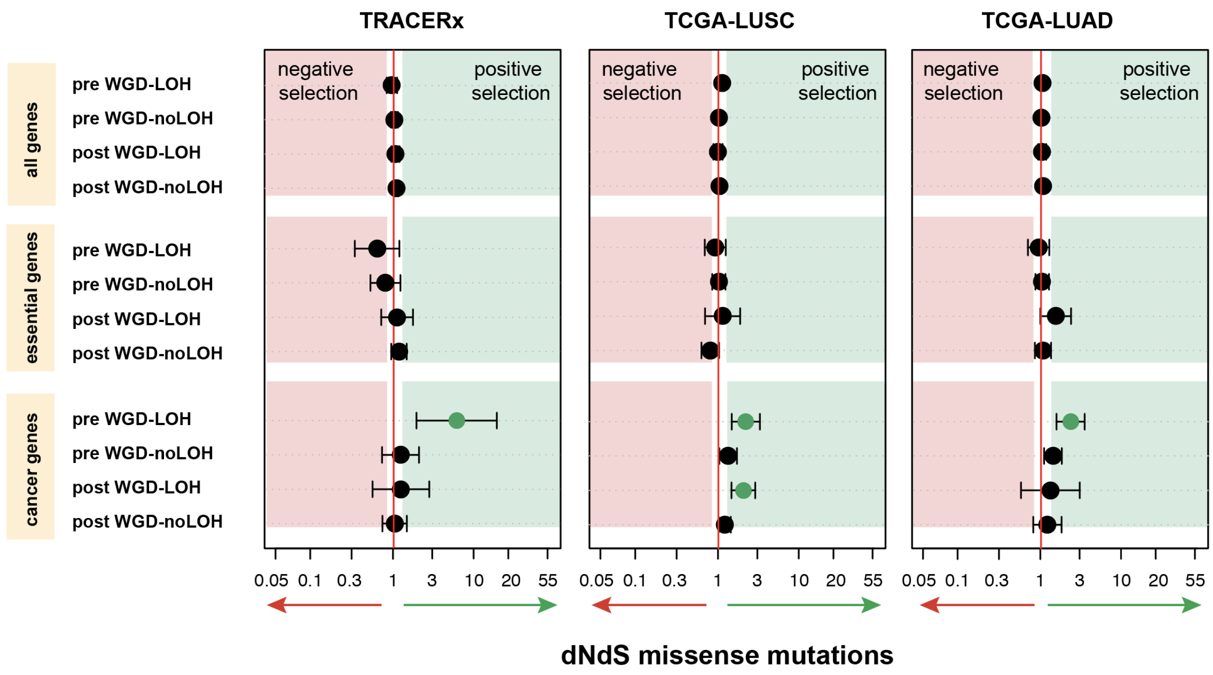

**Figure S4.** dNdS values for truncating mutations in WGD tumors calculated for different essential genes lists (Blomen et al., 2015; Hart et al., 2015 Wang et al., 2015) in TCGA-LUSC and TCGA-LUAD, grouped by LOH status and timing of the mutations.

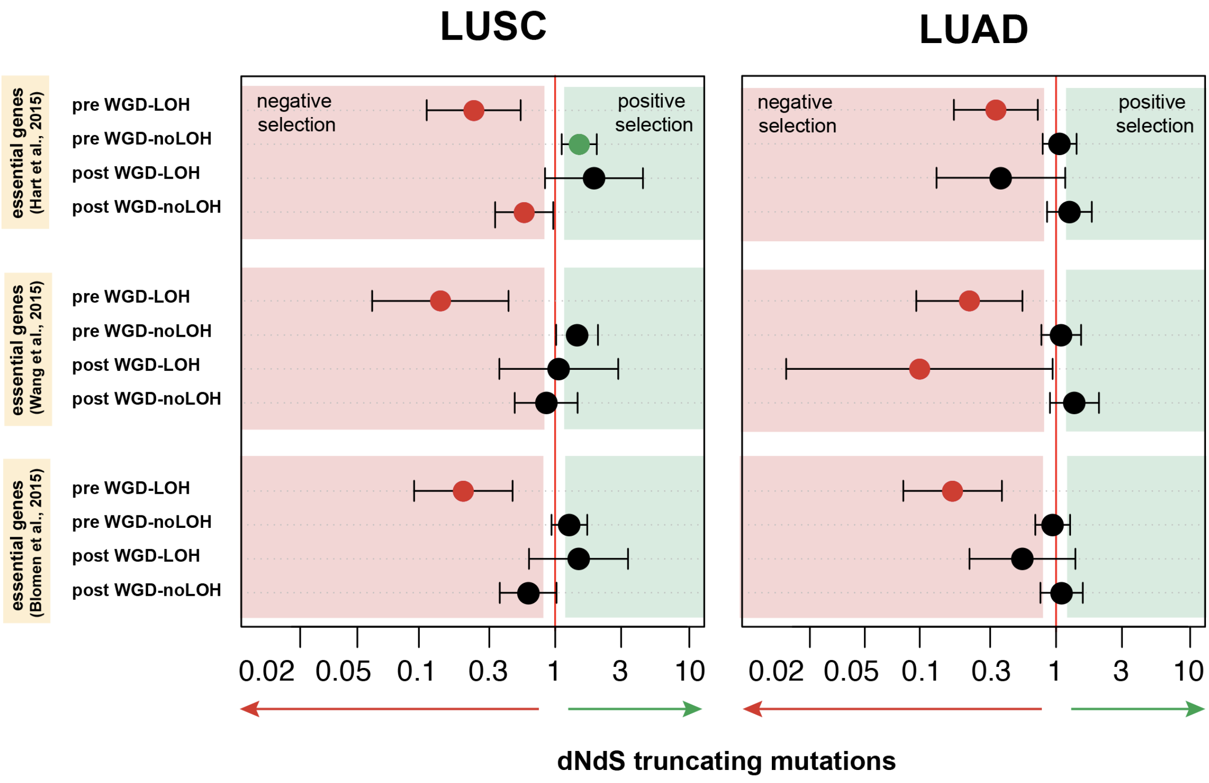

**Figure S5** Fractions of mean WGD and haploid LOH in non-WGD tumors across different cancer types from the TCGA cohort.

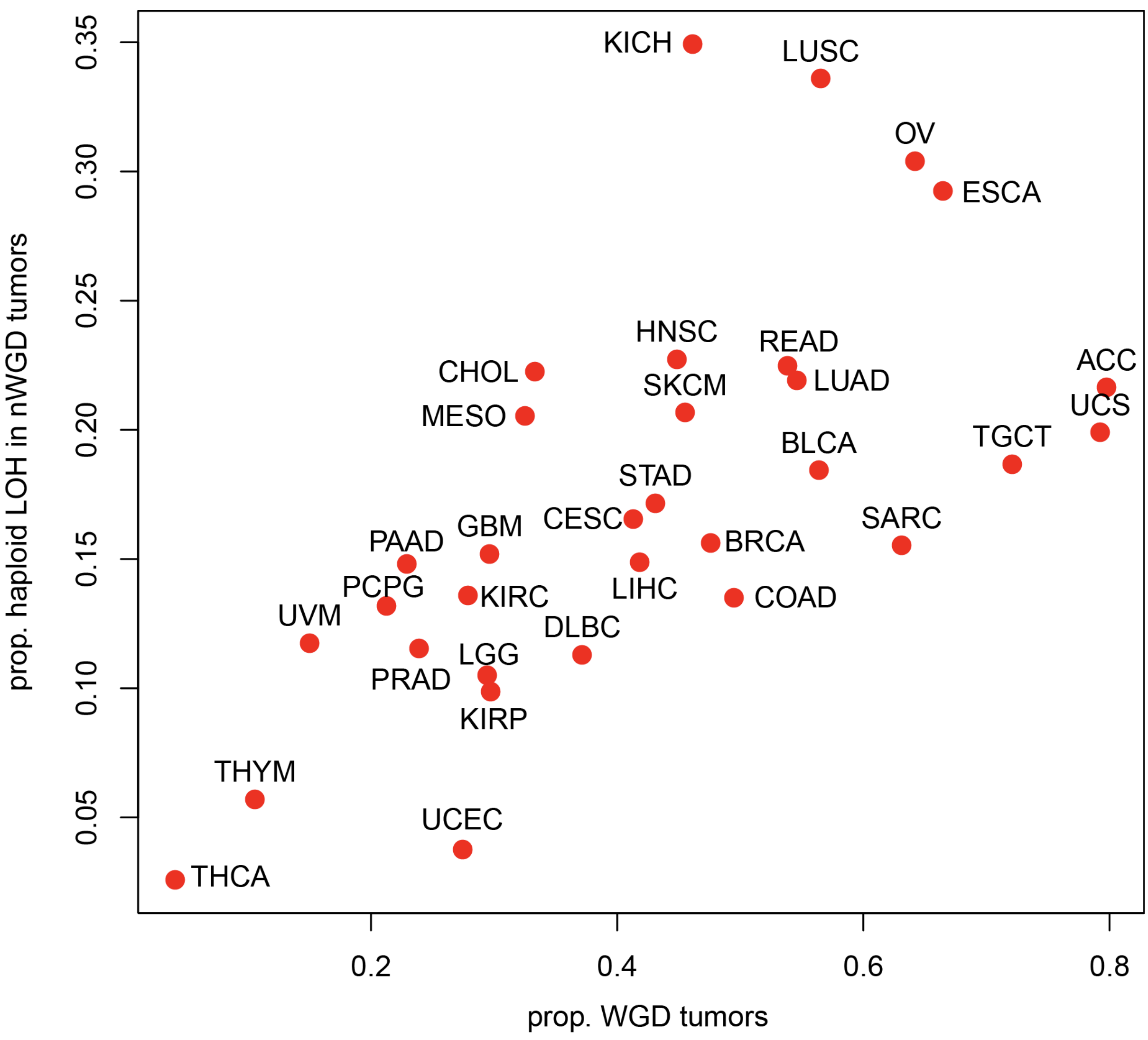

**Figure S6.** dNdS values for truncating mutations in WGD tumors calculated for essential genes, cancer genes, and all genes, grouped by LOH status and timing of the mutations in different cancer types from TCGA - MC3 calls (Ellrott et al., 2018).

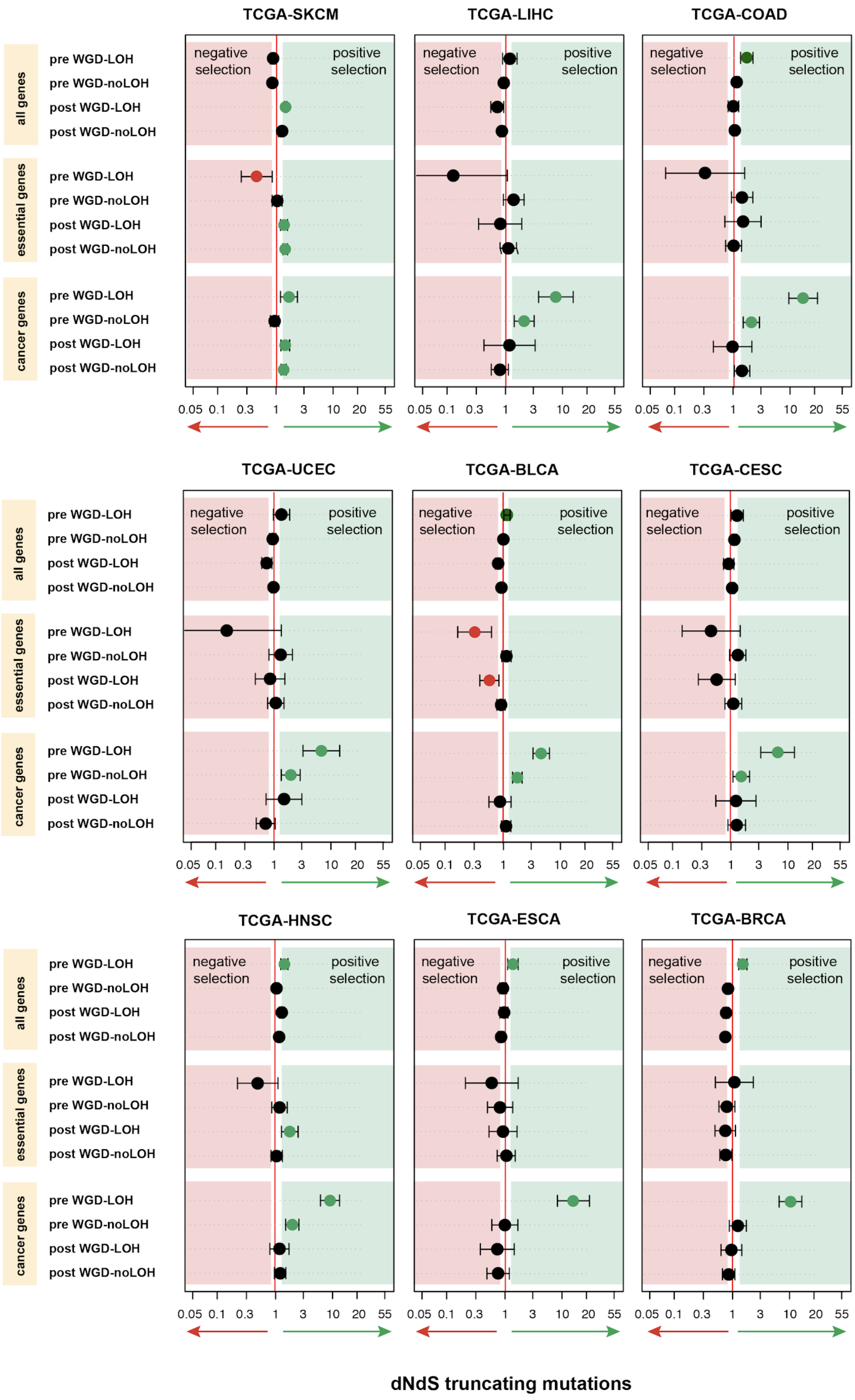
TCGA-SKCM
TCGA-LIHC
TCGA-COAD
TCGA-UCEC
TCGA-BLCA
TCGA-CESC
TCGA-HNSC
TCGA-ESCA
TCGA-BRCA
all genes
essential genes
cancer genes
pre WGD-LOH
pre WGD-noLOH
post WGD-LOH
post WGD-noLOH
negative selection
positive selection
0.05 0.1 0.3 1 3 10 20 55
dNdS truncating mutations

**Figure S7.** q-values for the significant genes (q-value < 0.05) identified by MutSigCV (Lawrence et al., 2013) in any of the 3 categories (early LOH, no LOH, all). The font color of the genes represents whether the gene is included in the COSMIC database (black=included, red=not-included).

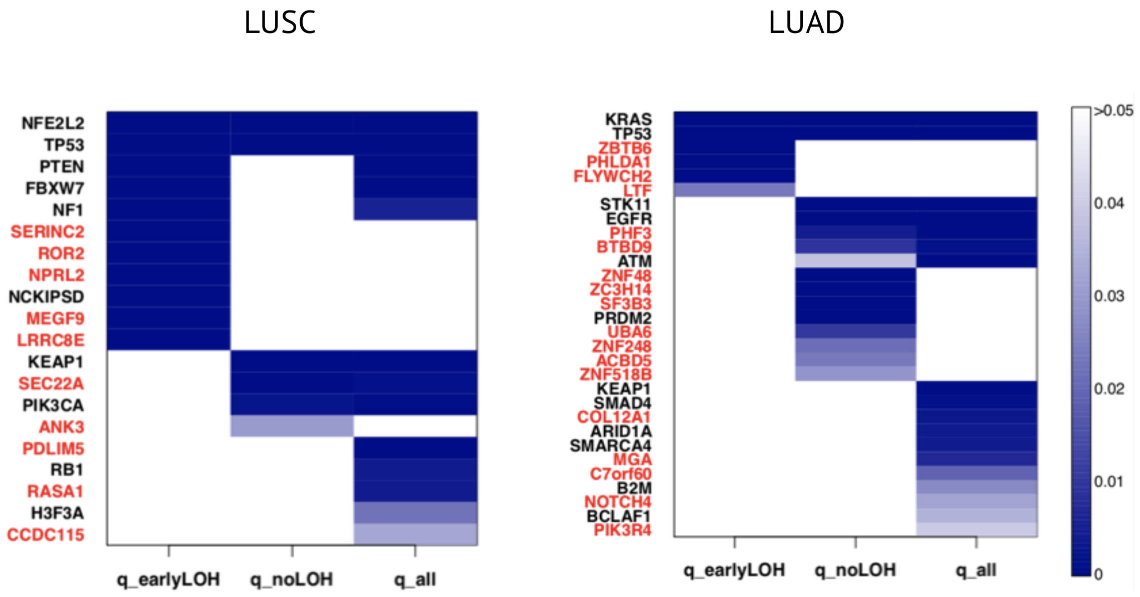
